# MINFLUX-nanoscopy of hNAIP/NLRC4 inflammasome activation in single human macrophages

**DOI:** 10.64898/2026.09.21.753160

**Authors:** Susanne Kulnik, Alexander Westerkamp-Carsten, Jonas Lübbe, Shuting Yin, Antonio Virgilio Failla, Martin Aepfelbacher

## Abstract

The biochemical and structural basis of NAIP/NLRC4 inflammasome activation is well understood. Far less well known are the spatiotemporal processes within cells that play a role in the activation of the NAIP/NLRC4 inflammasome. We used super-resolution imaging technology, including MINFLUX nanoscopy, to investigate the human hNAIP/hNLRC4 inflammasome in primary human macrophages following the phagocytosis of a specific *Yersinia enterocolitica* strain. hNAIP, which senses components of the bacterial type III secretion system (T3SS) and activates NLRC4, was recruited to bacteria-containing phagosomes after they were disrupted by the T3SS. Disruption of a single phagosome within a macrophage was sufficient to cause the accumulation of NLRC4 and NLRP3 (a sensor of another inflammasome), as well as the adapter protein ASC, in a dense condensate known as a speck. This subsequently led to plasma membrane permeation and pyroptosis. It appeared that, with increasing infection time, clusters containing NLRC4 and NLRP3/NLRC4 gradually acquired ASC for speck formation. MINFLUX-nanoscopy was able to visualise hNAIP at disrupted phagosomes with nanometre resolution, and also revealed the number and organisation of NLRC4 inflammasome discs within a speck, as well as their hNAIP content, for the first time. Surprisingly, the number of NLRC4 inflammasome discs varied considerably, ranging from 35 to 249 per speck. Also, only a small fraction of these seemed to contain hNAIP. Our results significantly improve our understanding of the process by which the hNAIP/NLRC4 inflammasome is activated in macrophages that have ingested bacteria containing a T3SS.

## Introduction

One of the immune system’s key functions is to detect pathogen-associated molecular patterns (PAMPs) and danger-associated molecular patterns (DAMPs) from infectious bacteria, fungi, viruses, and parasites. PAMPs are common structural components of pathogens, whereas DAMPs are a variety of molecules that signal infection. DAMPs are also produced as a consequence of the activity of microbial virulence factors/effectors, for example after they have disrupted crucial cellular signaling pathways. This process is known as effector-triggered immunity^1–4^.

In macrophages and other cell types, DAMPs and PAMPs induce the formation of heterologous multiprotein assemblies known as inflammasomes. These produce mature interleukin (IL) −1β and IL-18 and induce pyroptosis - a type of programmed cell death associated with inflammation - to stop the infection process^5,6^. Canonical inflammasomes consist of sensor proteins that contain a nucleotide-binding oligomerisation domain (NOD) and a leucine-rich repeat domain (LRR), as well as either a pyrin domain (PYD) or a caspase activation and recruitment domain (CARD). These different forms of inflammasome sensors are therefore named NLRPs or NLRCs, respectively^7,8^. Through their PYD, NLRPs induce the polymerisation of the adaptor protein ASC, which activates caspase-1. In comparison, NLRCs can directly bind to and activate caspase-1 via their CARD domain^5,9^. Active caspase-1 cleaves the IL-1β, IL-18 and gasdermin-D precursors, converting them into their mature forms. This leads to the assemble of gasdermin-D pores in the host cell plasma membrane, enabling the release of the mature IL-18 and IL-1β, and triggering pyroptosis^10–12^.

The enteropathogens *Yersinia enterocolitica* and *Y. pseudotuberculosis*, as well as the plague agent *Y. pestis*, interfere with inflammasomes at multiple levels^13^. Like many Gram-negative pathogens, pathogenic yersiniae use a type three secretion system (T3SS) to transport effector proteins into host cells^14^. The structural proteins of the T3SS, the needle and the inner rod, are sensed by NAIP (NAIP1,2,5 and 6 in mice and hNAIP in humans), which activates the NLRC4 inflammasome^15–18^. Furthermore, the translocation pore formed by the T3SS-secreted proteins YopB and YopD in the plasma membrane of the host cell enables entry of the T3SS effectors but also activates the NLRP3 inflammasome^19,20^. In addition, the antiphagocytic T3SS effectors YopE and YopT stimulate the pyrin inflammasome by inactivating Rho GTP-binding proteins^21–23^. Intriguingly, the yersiniae have developed an intricate network of mechanisms that effectively counteract activation of the multiple inflammasomes triggered by their T3SS structural and pore proteins and effectors. While all T3SS effectors work together to best inhibit inflammasome activation in human monocyte-derived macrophages (hMDMs)^24^, the YopP/YopJ and YopQ/YopK effectors (where the first Yop is the *Y. enterocolitica* homologue and the second is the *Y. pseudotuberculosis/Y. pestis* homologue) primarily block NLRP3- and NLRC4-inflammasome activities^19,20,24–26^. In comparison, the effector YopM primarily blocks the activation of the pyrin inflammasome induced by the effectors YopE and YopT^25,27,28^.

We recently showed that a small percentage of *Yersinia*-containing phagosomes rupture after integration of the bacterial translocation pore into the phagosomal membrane. Thereupon, galectin-3 and guanylate-binding protein 1 (GBP1) recruit to the now accessible bacteria^29^. GBP-1 can bind to bacterial LPS on the bacterial surface and, together with its isoforms, serves to recruit caspase-4, thereby triggering the formation of non-canonical inflammasomes^30–32^.

The current notion is that NAIP, the activator of the canonical NLRC4 inflammasome, binds to the needle and inner rod proteins of the T3SS in the cytoplasm of infected host cells and then initiates NLRC4 inflammasome assembly^16,17,33^. However, it is unclear how e.g., T3SS needle proteins would reach the cytoplasm if the bacteria are residing in phagosomes. We hypothesised that, as seen in the case of phagocytosed *Yersinia*, the bacterial T3SS components would also be accessible to NAIP after phagosome disruption. Therefore, we investigated in this study whether the activation of the NLRC4 inflammasome by the *Yersinia* T3SS in fact begins with the recruitment of NAIP to bacteria-containing disrupted phagosomes. Furthermore, although several in vitro studies have elucidated the structural details of NAIP/NLRC4 inflammasome activation in great detail^18,34^, the spatiotemporal characteristics of NAIP/NLRC4 activation within cells are not well understood. To address this deficit, we used an engineered *Yersinia* strain that produces the T3SS but does not translocate effectors, thereby achieving robust NLRC4 inflammasome activation in hMDMs. Using live-cell- and STED-microscopy as well as MINFLUX nanoscopy (with temporal resolution of minutes and spatial resolution of 5–10 nm), the entire process of NLRC4-inflammasome activation could be documented in single human macrophages. This process included phagosome disruption, hNAIP recruitment to the exposed bacterium, formation and maturation of different NLRC4-containing clusters over time, their coalescence into a single speck, and finally membrane permeation and pyroptosis.

## Results

### Activation of NLRC4 inflammasomes in single human macrophages after uptake of *Y. enterocolitica*

The aim of our study was to visualise at the single cell level and with high spatiotemporal resolution the process by which NLRC4 inflammasomes are formed after sensing of a bacterial type 3 secretion system (T3SS). The experiments were performed in living and fixed hMDMs infected with an engineered *Yersinia enterocolitica* strain. In humans, NLRC4 is activated by hNAIP, which detects the needle and inner rod proteins of the bacterial T3SS, whereupon hNAIP and NLRC4 form an inflammasome disc^15–17,34,35^. Notably, wild-type *Yersinia* spp. effectively block NLRC4 activation through the T3SS-mediated injection of effector proteins named Yops^20,24^. To achieve robust activation of the NLRC4-inflammasome, we therefore used an engineered *Yersinia* strain called WAC(pT3SS), which expresses the T3SS injection machinery including needle and inner rod proteins, but does not produce and translocate the Yops^36^. Macrophages infected with WAC(pT3SS) exhibited a time-dependent formation of ASC specks, which are single sites in the cell where inflammasomes cluster (Fig. 1A)^37–39^. Coimmunostaining demonstrated that the ASC specks can contain NLRC4, its activator hNAIP, caspase-1 and NLRP3 (Fig. 1B). The infected macrophages exhibited time-dependent permeation of the plasma membrane and the formation of membrane blebs, which are characteristic of pyroptotic cell death. They also secreted significant amounts of IL-1β (Fig. 1C-E). Movies of WAC(pT3SS)-infected macrophages expressing green fluorescent protein (GFP)-ASC demonstrated that ASC speck formation precedes plasma membrane permeation, indicating a causal relationship (Fig. 1F). Neither the plasmid cured avirulent strain WAC nor the wild type strain WA-314 induced any of these hallmarks of pyroptosis, as they either do not produce a T3SS (WAC) or block the T3SS-induced activation of NLRC4 through the Yops (WA-314) (Fig. 1A, C, E)^20^. We conclude that *Yersinia* WAC(pT3SS) induces functionally active NLRC4/hNAIP inflammasomes after ingestion by primary human macrophages, leading to pyroptosis. The presence of NLRP3 in the ASC specks suggests that other inflammasomes may also contribute to the WAC(pT3SS) effect on pyroptosis.

**Figure 1:**
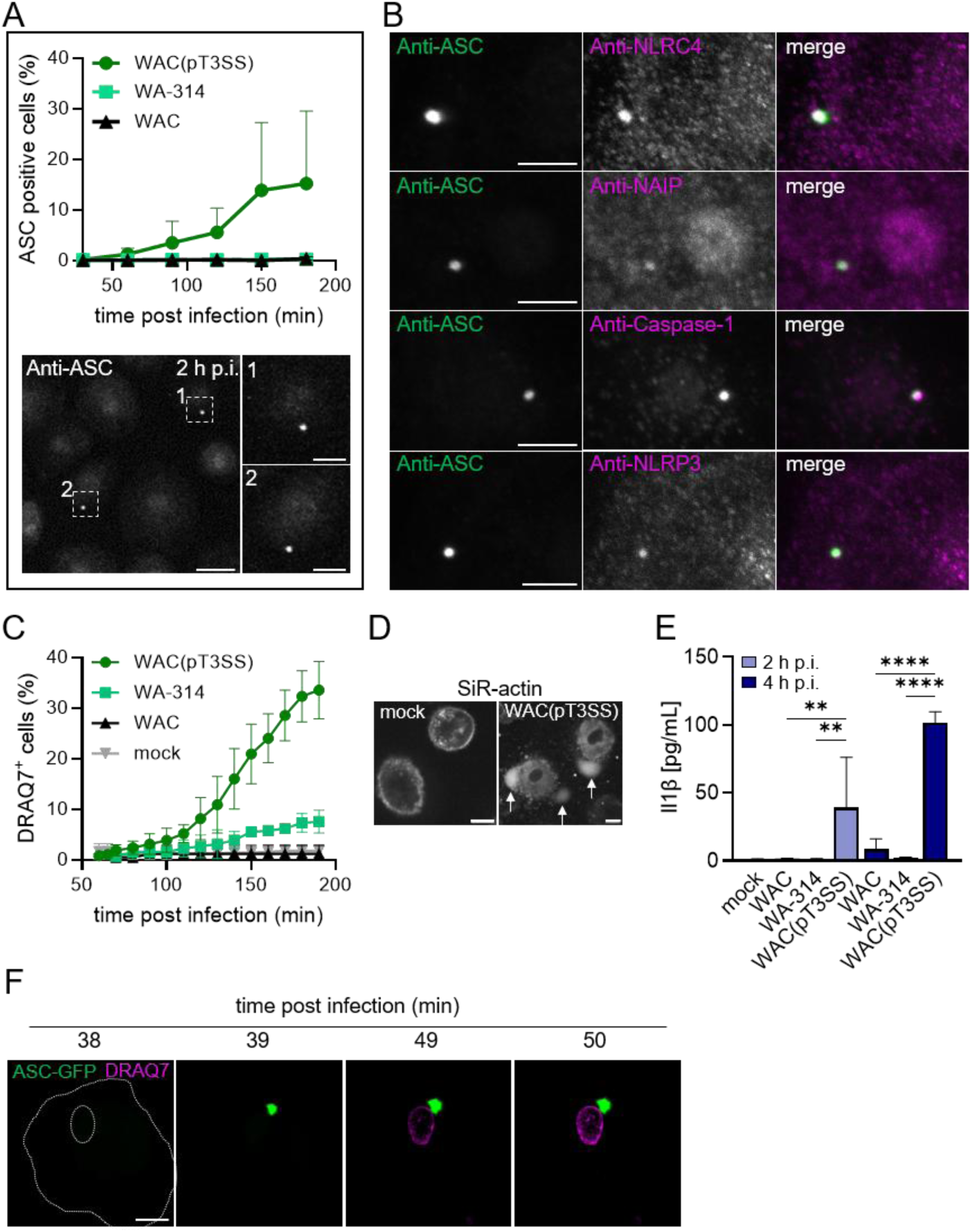
*Y. enterocolitica* WAC(pT3SS) activates NLRC4 inflammasomes in hMDMs. **A) Time course of ASC speck formation.** Macrophages were infected with *Y. enterocolitica* WAC, WA-314 or WAC(pT3SS) for 30 to 180 min and immunostained with primary α-ASC and fluorescently-labelled secondary antibody. Data of ASC speck positive cells are the mean ± SD of at least three biological replicates. Images depict a representative overview and blow-ups of individual ASC specks after 120 min of infection. Scale bar: 20 µm (overview) and 10 µm (blow-up**). B) Colocalisation of ASC and other inflammasome components within specks.** hMDMs were infected with *Y. enterocolitica* WAC(pT3SS) for 120 min and immunostained with primary antibodies and fluorescently labelled secondary antibodies to visualise ASC (green) and either one of the inflammasome components NLRP3, caspase-1, NLRC4 or hNAIP (magenta). Representative images are shown. Scale bar: 5 µm. **C) Plasma membrane permeation in infected macrophages**. DRAQ7^TM^ fluorescent, membrane impermeable dye was added during infection with indicated strains for the indicated times and DRAQ7 positive cells were detected with spinning-disk live-cell microscopy. Depicted are the mean ± SD of at least three biological replicates. **D) Membrane blebbing in infected macrophages.** Macrophages were not infected (mock) or infected with WAC(pT3SS) for 2 h and stained with SiR-actin. Arrowheads point to membrane blebs. Scale bar: 10 µm. **E) IL-1β release in infected macrophages.** Macrophages were not infected (mock) or infected with WAC, WA-314 or WAC(pT3SS) for 2 h or 4 h and IL-1β concentration (pg/mL) in the supernatant was measured by ELISA. Each bar represents the mean ± SD of at least three biological replicates (** p < 0.01, **** p < 0.0001; ordinary one-way ANOVA). **F) ASC speck formation is followed by cell death in macrophages.** ASC-GFP (green) expressing hMDMs were infected with WAC(pT3SS) in the presence of DRAQ7^TM^ dye (magenta) and ASC specks and DRAQ7 positive nuclei were imaged at indicated time points with spinning disk microscopy. Scale bar: 10 µm. All infections were performed with an MOI of 100.

### The NLRC4 activator hNAIP is recruited to disrupted *Yersinia* containing phagosomes

The current notion is that hNAIP binds to the needle and inner rod proteins of the T3SS in the cytoplasm of infected host cells. However, it is unclear how e.g., needle proteins would reach the cytoplasm^16,17,33^. We recently showed that a small percentage of *Yersinia*-containing phagosomes rupture after integration of the T3SS translocation pore into the phagosome membrane^29^. Against this background, we investigated whether hNAIP is recruited to *Yersinia*-containing ruptured phagosomes. To this end, we co-expressed the specific marker for disrupted phagosomes, mScarlet-galectin-3^40^, and eGFP-hNAIP in macrophages. We then infected the macrophages with WAC(pT3SS) and examined the colocalisation of the two markers, either over time using live-cell imaging or after fixing the cells. eGFP-hNAIP was regularly detected at mScarlet-galectin-3 positive phagosome remnants in fixed cells (Fig. 2A, S1D). In the fixed macrophages we also detected dot like accumulations of endogenous hNAIP at the bacteria (Fig. 2B; S1C). Live-cell imaging indicated that mScarlet-galectin-3 clearly accumulated before eGFP-hNAIP at the phagosomes (Fig. 2C). In addition, we tested by live-cell imaging whether the hNAIP binding partner NLRC4 is also recruited to disrupted phagosomes. In movies of infected macrophages expressing mScarlet-galectin-3 and eGFP-NLRC4, it appeared that neither before nor at any time point after phagosome disruption eGFP-NLRC4 was present at the phagosome (Fig. 2D). In contrast, only after mScarlet-galectin-3 recruitment, eGFP-NLRC4 started to accumulate in specks at some distance to the disrupted phagosome (Fig. 2D). Altogether, we propose that hNAIP senses *Yersinia* T3SS proteins at disrupted phagosomes, where it is activated and then moves to the specks at some distant cellular localisation. There or on its way there it stimulates NLRC4.

**Figure 2:**
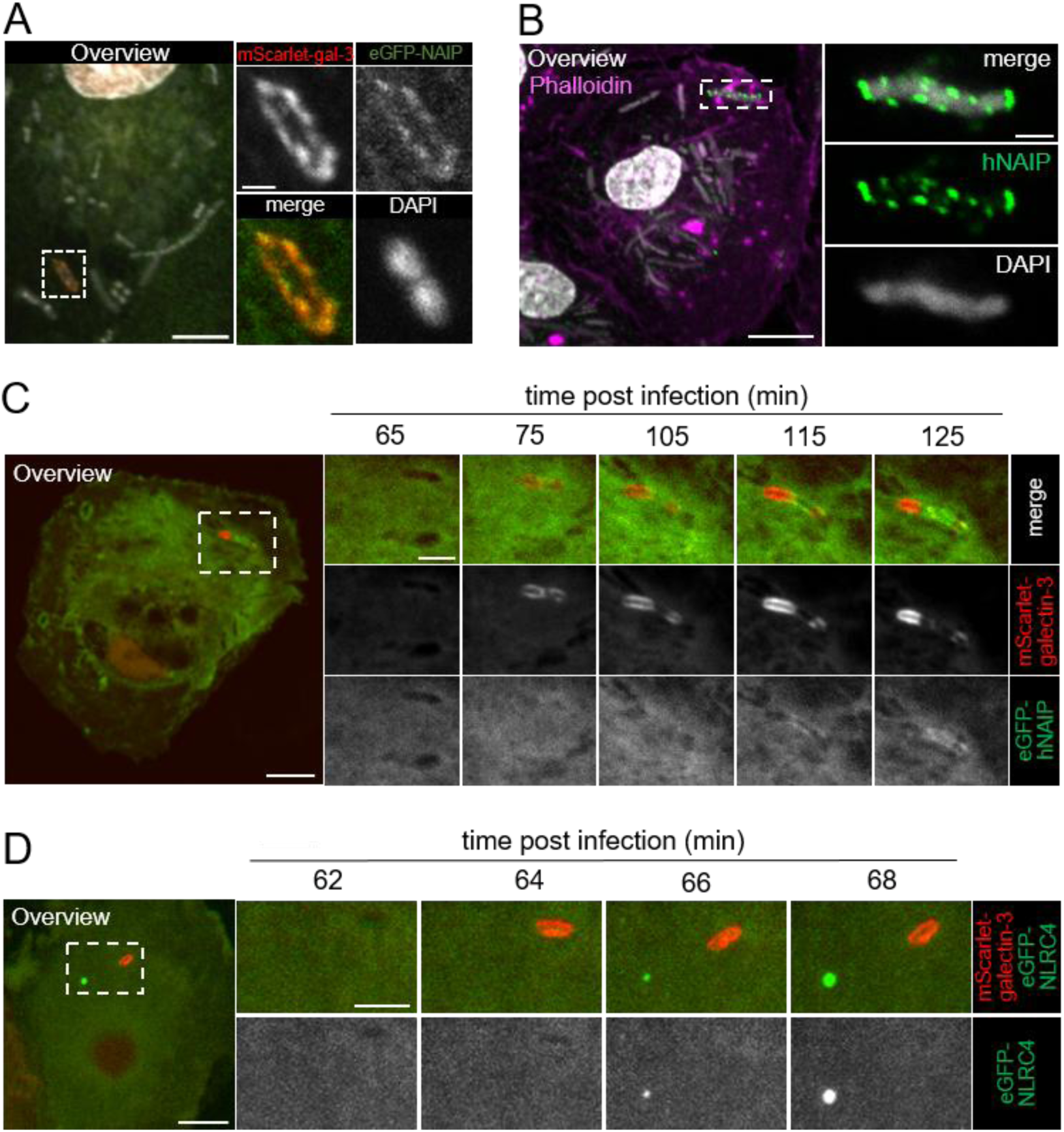
hNAIP is recruited to disrupted phagosomes in *Y. enterocolitica* infected macrophages. **A) Colocalization of mScarlet-galectin-3 and hNAIP at a bacterium-containing phagosome.** hMDMs expressing mScarlet-galectin-3 (red) and eGFP-hNAIP (green) were infected with WAC(pT3SS) at an MOI of 100 for 1 h, stained with DAPI (white) and fixed. Confocal images show a representative overview and the enlargement of the indicated area is shown as red, green and white channels and merge depicted separately. Scale bars: overview: 5 µm, blow-up: 1 µm. **B) Endogenous hNAIP localizes to the bacteria.** hMDMs were infected with WAC(pT3SS) at an MOI of 100 for 3 h, and immunostained for hNAIP (green) and additionally stained with DAPI (white) and phalloidin (magenta), to visualise bacterial/cellular DNA and the cellular actin-cytoskeleton, respectively. A representative overview and the blow-up of the indicated area with merge, green and white channels shown separately are depicted. Scale bars: overview: 10 µm, blow-up: 2 µm. **C) Recruitment of eGFP-hNAIP to a ruptured phagosome**. hMDMs were transfected and infected as in A. Stills from a representative movie at indicated time points recorded with live-cell spinning disk microscopy are shown. Scale bars: overview: 10 µm, blow-up: 2 µm. **D) NLRC4 accumulates at the speck over time but not at the ruptured phagosome.** hMDMs expressing mScarlet-galectin-3 (red) and eGFP-NLRC4 (green) were infected with WAC(pT3SS) at an MOI of 100 and recorded with live-cell spinning disk microscopy. An overview and representative stills of the boxed area at indicated timepoints post infection are shown. Scale bars: overview: 10 µm, blow-up: 5 µm.

### The disruption of one phagosome drives human macrophages into pyroptosis

Towards our goal to investigate the spatio-temporal characteristics of NLRC4 inflammasome activation at the single cell level, we employed movies of WAC(pT3SS)-infected single macrophages expressing mScarlet-galectin-3 and ASC-GFP. With these we analysed: i) how long it takes for ASC specks to form after they are triggered at the phagosome; ii) whether the distance in the cell between the disrupted phagosome and the ASC speck correlates with the time it takes for the ASC speck to form; and iii) how many disrupted phagosomes are needed to trigger the formation of an ASC speck. The 22 analysed macrophages from three different donors showed considerable heterogeneity in terms of the time after infection at which phagosomes disrupted (from 40 min to 150 min) and the time after phagosome disruption at which ASC specks formed (from 0 min to 71 min) (Fig. 3A). Furthermore, the distance between the disrupted phagosome and the resulting ASC speck in the macrophages varied between 0 µm and 28 µm, but this distance did not correlate with the time it took for the specks to form (Fig. 3B). It is worth noting that in almost 90% of cases where a speck formed in a macrophage, this was triggered by one (59%) or two (30%) disrupted phagosomes (Fig. 3C). These results not only reveal significant differences in the timing of phagosome disruption and inflammasome activation in single human macrophages after they have phagocytosed *Yersinia*. They also provide compelling evidence that an infected macrophage can be driven into pyroptosis by just one disrupted phagosome.

**Figure 3:**
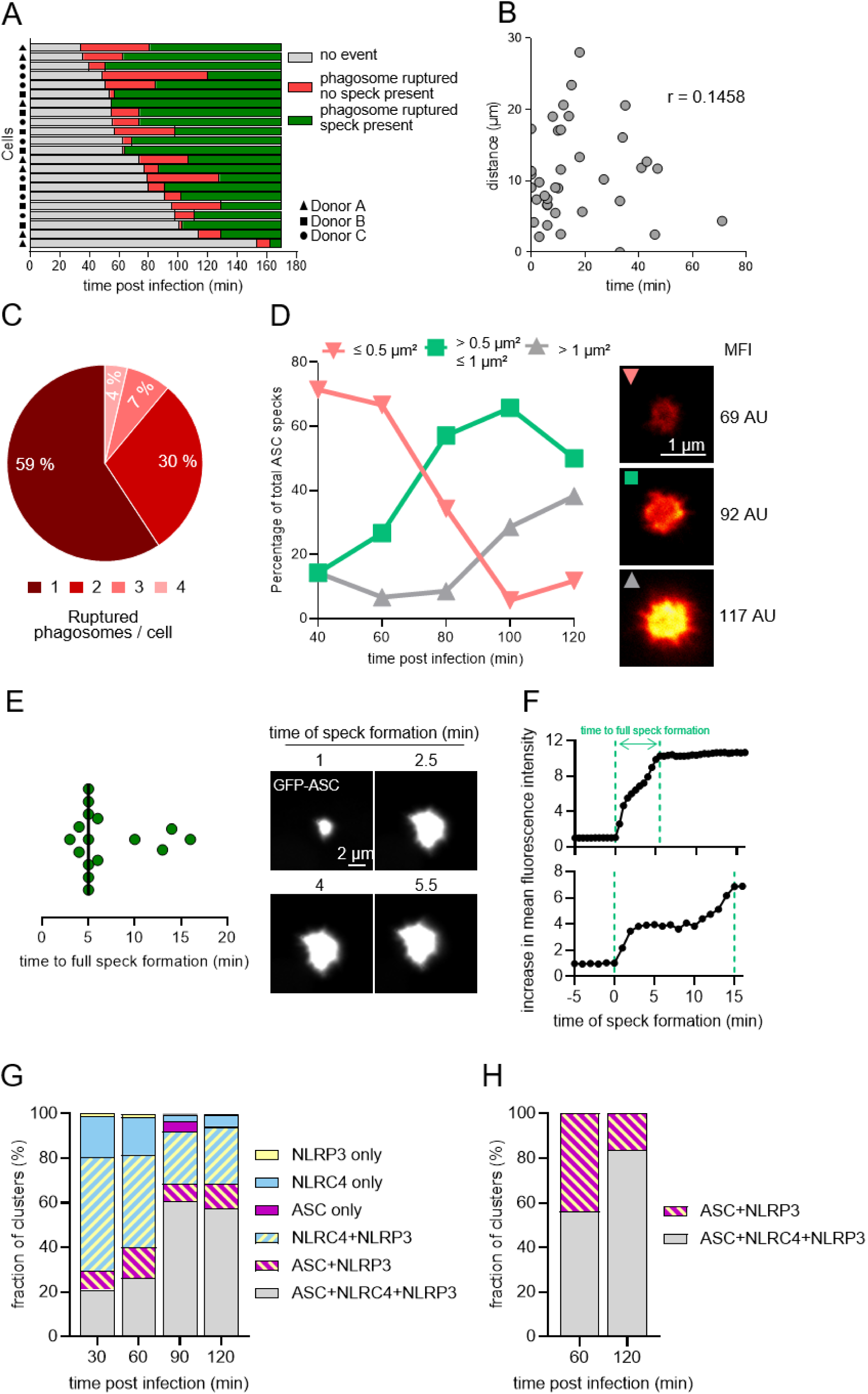
Dynamics and variability of ASC speck formation and composition in infected macrophages. **A) Kinetics of phagosome rupture (red) and ASC-GFP speck formation (green) in infected macrophages.** hMDMs from three different donors expressing mScarlet-galectin-3 and ASC-GFP were infected with WAC(pT3SS) at an MOI of 100 and imaged with live-cell spinning disc microscopy. The times of phagosome rupture and ASC-GFP speck formation were recorded and depicted for each single cell starting with the cell showing the fastest phagosome rupture. **B) Correlation** between time it took from phagosome rupture to speck formation and distance from ruptured phagosome to the formed speck in the same cell. r = 0.1458. Data are from A. **C) Number of ruptured phagosomes per macrophage.** The pie diagram shows that 59 % of the cells displayed one ruptured phagosome before ASC-GFP speck formation, 30 % displayed two, 7 % displayed three and 4 % displayed four. Data are from A. **D) ASC specks increase in size during macrophage infection.** hMDMs were infected with WAC(pT3SS) at an MOI of 300 for indicated times, immunostained for endogenous ASC specks and imaged with 2D-STED super-resolution microscopy. Sizes of ASC specks were attributed to three categories: Red: smaller than 0.5 µm²; Green: bigger than 0.5 µm² and smaller than 1 µm²; Grey: bigger than 1 µm². Number of ASC specks measured for each time point: 40 min: 7; 60 min: 30; 80 min: 35; 100 min: 35 min; 120 min: 34. Specks representative for each size and the respective MFI (mean fluorescence intensity) are depicted. Scale bar: 1 µm. **E) Time to full ASC speck formation in single infected macrophages**. ASC-GFP expressing hMDMs were infected with WAC(pT3SS) at an MOI of 100 and imaged with live-cell spinning disk microscopy. The time it took from first visible accumulation of ASC-GFP until the ASC-GFP fluorescence intensity plateau was reached in individual specks was determined (see F for scheme). Each dot represents one of 16 analysed ASC specks. The vertical line depicts the median. **F) Time courses of the mean ASC-GFP fluorescence intensities** in two representative developing specks from E are depicted. The time between the dashed green lines represents time for full speck formation. **G) The composition of inflammasome clusters changes over time in infected macrophages.** hMDMs were infected with WAC(pT3SS) at an MOI of 100 for indicated times and immunostained for clusters of endogenous ASC, NLRP3 and NLRC4. Data are mean of 3 biological replicates of at least 73 clusters per timepoint. **H) Dynamic composition of ASC containing clusters between 60 and 120 min of infection.** Shown are the percentages of ASC + NLRP3 and ASC + NLRP3 + NLRC4 containing clusters from G at the 60 and 120 min timepoints.

### Size and composition of NLRC4 clusters in infected macrophages develop dynamically over time

In order to get a better idea of the stages involved in the formation of ASC specks, we measured their size and fluorescence intensity in macrophages infected with WAC(pT3SS) over a period of 120 minutes. At 40 min post infection, the percentage of small ASC specks (area <0.5 μm²) was at around 75 % whereas medium size (area 0.5 to 1 μm²) and large (area > 1 μm²) specks where below 20 %. Until 120 min of infection, the percentage of small specks dropped to around 15%, whereas the medium size and large specks rose to around 45 % and 40 %, respectively (Fig. 3D). These data provide a first hint that ASC specks develop gradually in an infected macrophage population. Because the data are based on snapshots in fixed macrophages, they do not provide an insight into the time it takes for an individual ASC speck to develop. To answer this question, the increase in the mean intensity of the developing ASC-GFP specks from the cells in Fig. 3A was measured. The median time it took an ASC-GFP speck to fully form was found to be 5 min (Fig. 3E). Whilst around three fourth of specks formed after around 5 min, the remaining fourth took between 10 and 16 min to develop fully (Fig. 3E). The slow-developing specks showed a biphasic increase in GFP-ASC fluorescence (Fig. 3F). We therefore speculate that the formation of ASC specks follows a standard programme that can be halted and resumed in some cases, resulting in a biphasic increase. Due to its ease of detection, ASC (mainly in the form of ASC-GFP) has been widely used as a marker for the clustering of inflammasome components within a cell. The rather large ASC-GFP clusters that form were referred to as ASC specks^41^. To gain a more unbiased insight into the clustering of inflammasome components, we performed triple staining of endogenous ASC, NLRC4 and NLRP3 in macrophages over a period of 120 min of infection. The results showed that clusters comprising almost every possible combination of the three components were found, albeit in varying proportions at different points in infection time (only ASC/NLRC4 was missing) (Fig. 3G). During the observation period of 30 to 120 minutes, the following changes occurred: i) Clusters containing ASC/NLRC4/NLRP3 increased from 20% to approximately 60% of all detected clusters; ii) Clusters containing NLRC4/NLRP3 decreased from approximately 50% to 20%; iii) Clusters containing only NLRC4 dropped from 10% to less than 5%; iv) Clusters containing ASC/NLRP3 varied between 10% and 15%. Clusters containing only NLRP3 or ASC were present only in minor amounts (Fig. 3G). These data indicate that up to 120 minutes after infection, the proportion of ASC/NLRC4/NLRP3 clusters increases considerably, at the expense of NLRC4/NLRP3 and NLRC4 clusters. Because we did not observe ASC/NLRC4 clusters, these findings suggest that ASC is recruited to pre-existing NLRP3/NLRC4 clusters. When considered alongside the small vs. medium/large ASC-containing clusters that prevail 60 and 120 minutes after infection, respectively (see Fig. 3D), it also becomes clear that the smaller ASC specks seen after 60 min are more likely to only contain NLRP3, whereas the medium/large ASC specks at 120 min contain NLRP3 and NLRC4 (see Fig. 3H).

### MINFLUX-nanoscopy visualises single NLRC4 inflammasomes and hNAIP within ASC-specks

In the next step, we sought to examine the architecture of mature specks (formed after 120 min of infection) with regard to the arrangements of ASC, NLRC4 and hNAIP. For this, we employed 3D-STED super-resolution microscopy and visualised specks in eGFP-NLRC4-expressing macrophages that were stained for endogenous ASC and hNAIP. ASC and NLRC4 appeared as ring-like structures in the xy, xz and yz planes of the specks (Fig. 4A). 3D reconstructions confirmed that these structures form a hollow sphere (Movie S2). Surprisingly, hNAIP showed a patchy localisation at the NLRC4 spheres (Fig. 4A). Further image analysis suggested that ASC and NLCR4 form an outer and an inner shell of a speck (Fig. 4B-D).

**Figure 4:**
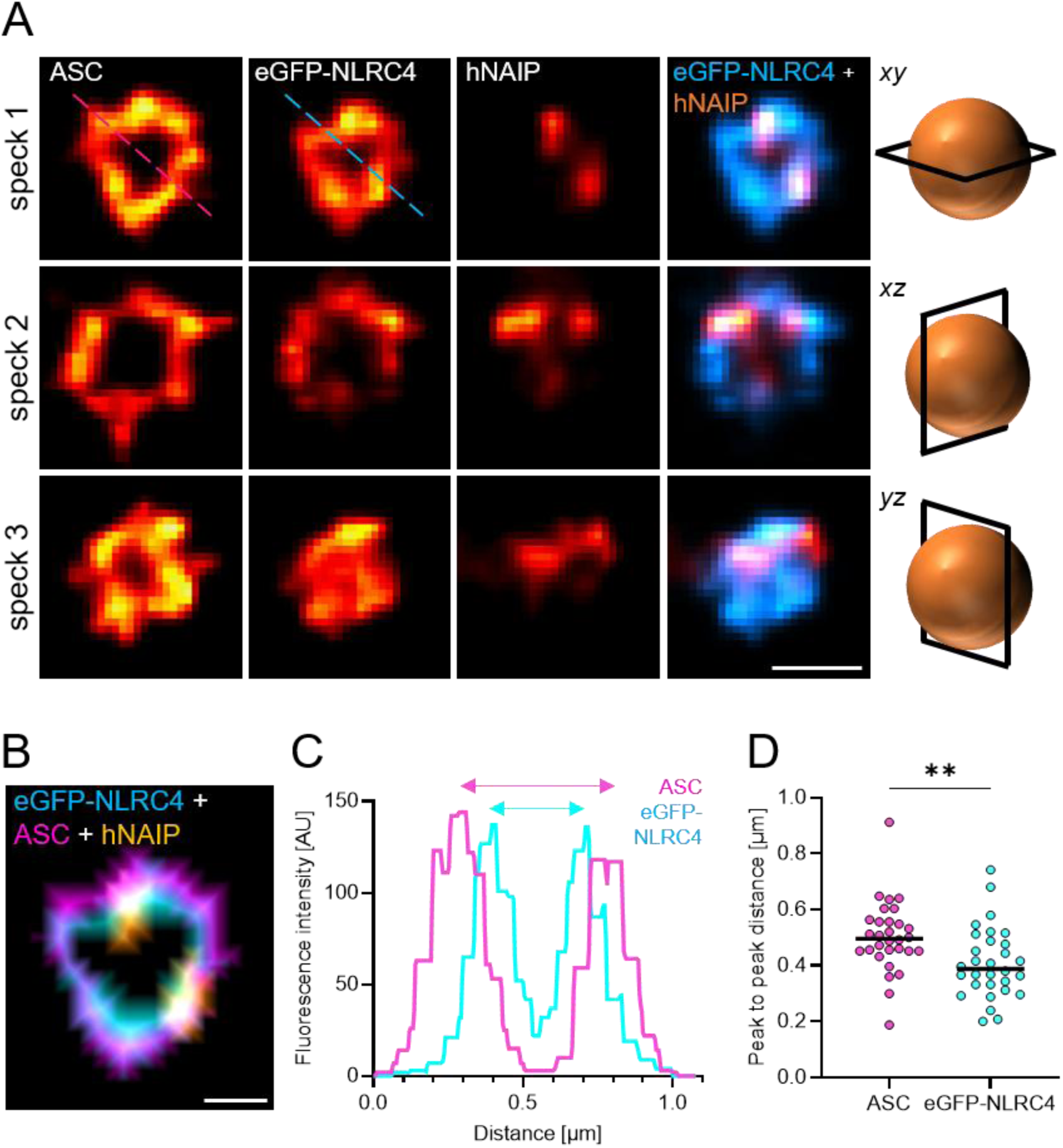
Suborganisation of ASC, NLRC4 and hNAIP in mature inflammasome clusters/specks. **A) Localisation of ASC, eGFP-NLRC4 and hNAIP in specks of infected macrophages.** hMDMs expressing eGFP-NLRC4 were infected with WAC(pT3SS) at an MOI of 100 for 120 min and immunostained for endogenous ASC and hNAIP as well as eGFP-NLRC4 (Methods). Each component is shown separately in three independent specks (1,2,3), each with a different orientation (xy, xz and yz). Merges of eGFP-NLRC4 (cyan) and hNAIP (orange) are also depicted (right column of images). The dashed lines in the ASC (magenta) and eGFP-NLRC4 channels (cyan) indicate the plot lines that were used for generating Fig. 4C. Scale bar: 500 nm. B) Representative distance map of the distribution of ASC (magenta), eGFP-NLRC4 (cyan) and hNAIP (yellow) within an ASC speck based on speck 1 in A. The distance map was created by processing binary images of ASC, eGFP-NLRC4 and hNAIP channels in Fiji. Scale bar: 500 nm. C) Representative plot profiles of the fluorescence intensities of ASC (magenta) and eGFP-NLRC4 (cyan) along the dashed lines indicated in A. The arrows exemplarily indicate the distances taken for peak to peak measurements in D. D) **Peak to peak distances of ASC and eGFP-NLRC4** along line profiles as depicted in C. Each dot represents one peak to peak distance. For each speck, two measurements per speck component and orientation (*xy*, *xz*, *yz)* using two different line profiles were performed. In total, ASC and eGFP-NLRC4 within five different specks were evaluated. Welch’s t test p-value: 0.0065.

Although STED microscopy was well suited for gaining insight into the overall arrangements of ASC, NLRC4 and hNAIP within a speck, its resolution is insufficient for visualising individual inflammasome disks. To achieve this, we employed MINFLUX nanoscopy, a technology that can localise individual fluorescent molecules emitting a relatively small number of photons. This enables the localisation of fluorescent dyes with a precision of a few nanometres (1–10 nm)^42–45^. In the first step we imaged endogenous NLRC4 and ASC in mature specks using primary antibodies and secondary nanobodies and exchange-PAINT. The obtained MINFLUX images revealed numerous agglomerates of ASC and NLRC4 molecules whereby the number of ASC agglomerates far exceeded that of the others (Fig. 5A). This is probably because, as demonstrated in vitro, NLRC4- and NLRP3-discs can stimulate the formation of substantial ASC filaments comprising a large quantity of molecules^9,46,47^. When the NLRC4 agglomerates were analysed in detail, it was found that their maximal diameter was around 45 nm (Fig. 5B).

**Figure 5:**
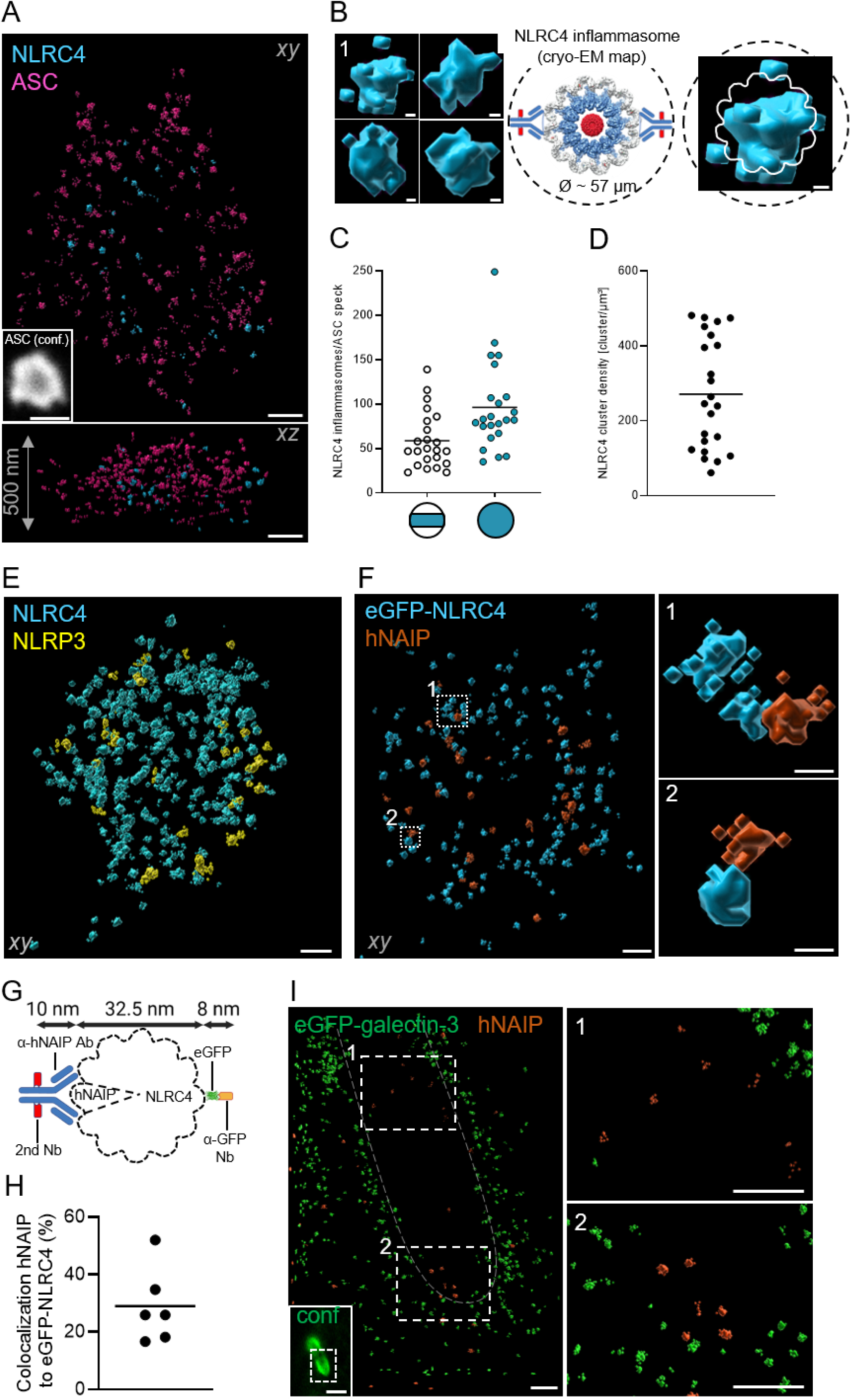
3D-MINFLUX nanoscopy resolves NLRC4 inflammasomes in specks and hNAIP at disrupted phagosomes at the nanometre scale. hMDMs were infected with *Y. enterocolitica* strain WAC(pT3SS) using an MOI of 100 for 2 h, if not indicated otherwise. Immunostainings for MINFLUX nanoscopy were performed using primary antibodies against indicated components and secondary nanobodies/antibodies (Methods). eGFP-tagged proteins were stained using an α-GFP nanobody. Altogether this enabled (consecutive exchange-) DNA-PAINT using imager strands coupled to Abberior DNA-PAINT 660 or Atto655. All MINFLUX localisations are depicted using the Imaris surface function (surface detail: 2 nm). **A) Nanoscale localisations of ASC (magenta) and NLRC4 (cyan) within an ASC speck.** ASC was additionally stained using a fluorescently labelled secondary α-mouse antibody for the confocal mode. Top: MINFLUX image: xy-view; Bottom: MINFLUX image: xz-view. The z-range was 500 nm. Inset (bottom left): Confocal image of ASC in the depicted speck. Scalebars: MINFLUX images: 200 nm; Confocal image: 1 µm. Infection was performed with an MOI of 300. **B) Comparison of single NLRC4 clusters to published cryo-EM data of the in vitro assembled NLRC4 inflammasome**. Left: blow-up of single NLRC4 clusters from A; Published cryo-EM map of the huNLRC4 inflammasome from Matico et al. ^34^ including antibody sizes and an overlay of the cyro-EM outline on the MINFLUX data of inflammasome 1. Scale bars: 5 nm. **C) Number of NLRC4 inflammasomes per ASC speck.** NLRC4 surfaces were counted into one cluster if their centre of masses had a maximal distance of 20 nm (Methods). Dots at the left circle with cyan stripe depict the number of NLRC4 clusters actually measured in the images because of the limited z-range. The dots at the fully cyan-coloured circle depict the calculated numbers of NLRC4 clusters within a whole speck assuming that specks roughly have a spherical shape (compare Fig. 4A). Each dot represents the number of NLRC4 clusters within one ASC speck. Horizontal lines depict the mean. **D) Density of NLRC4 clusters per µm³ of ASC specks**. Each dot represents the cluster density within one ASC speck. The horizontal line depicts the mean**. E) Nanoscale localisations of NLRC4 (cyan) and NLRP3 (yellow) within a putative speck**. hMDMs were infected with WAC(pT3SS) using an MOI of 300 for 2 h. NLRC4 was additionally stained using a fluorescently labelled secondary α-rabbit antibody for the confocal mode. Scalebar: 100 nm; xy-view. **F) Nanoscale localisations of NLRC4 (cyan) and hNAIP (red) within a putative speck.** Dashed boxes indicate region of blow-ups on the right. Scale bar: 100 nm; xy-view. Blow-ups: Scale bar: 20 nm. **G) Scheme for the estimation of hNAIP and eGFP-NLRC4 colocalisation.** The maximal distance hNAIP and eGFP-NLRC4 can maximally have within one inflammasome disk, including the size of the labels that were used for hNAIP (primary antibody and secondary nanobody; ∼ 10 nm), the size of a single inflammasome disk (32.5 nm;^34^), the sizes of eGFP fused to NLRC4 and the GFP-binding nanobody (8 nm) together with the approximately achieved localization precision (∼ 5 nm) is ∼ 55 nm. Created in BioRender. Carsten, A. (2026) https://BioRender.com/58x7puk **H) Percentage of hNAIP localisations colocalising with eGFP-NLRC4 clusters.** Surfaces were counted into one agglomeration if their centre of masses have a maximal distance of 20 nm. hNAIP and eGFP-NLRC4 agglomerations were considered as colocalising if their center of masses are within a distance of 55 nm (according to F). Each dot represents the percentage of hNAIP localisations that colocalised with eGFP-NLRC4 within one putative speck. The horizontal line depicts the mean. **I) hNAIP at a bacterium-containing disrupted phagosome in an infected macrophage.** hNAIP was additionally stained using a fluorescently labelled secondary α-rabbit antibody for localising hNAIP at ruptured phagosomes using the confocal mode. Ruptured phagosome indicated by the eGFP-galectin-3 fluorescence signal in confocal microscopy (inset in lower left corner). Dashed box indicates area of depicted MINFLUX image. Scale bar: 2 µm. MINFLUX: eGFP-galectin-3 (green) and hNAIP (red) using 3D-MINFLUX nanoscopy. Dashed boxes indicate blow-ups depicted on the right. Scale bar: 200 nm. Right: Blow-ups of hNAIP and eGFP-galectin-3. Scale bar: 200 nm.

The in vitro diameter of an NLRC4 disc has been determined to be approximately 32 nm^34,48^ Therefore, the maximal diameter of the NLRC4 agglomerates of 45 nm determined by MINFLUX nanoscopy is consistent with them representing individual inflammasome discs considering offset of the labels and localization precision^49^. Now that we were for the first time able to visualise single NLRC4 discs, we could determine their number in individual specks. It turned out that this number varied considerably, ranging from 35 to 249 per speck (Fig. 5C). Due to the limited z-range of the current MINFLUX application, we can infer that the actual number of NLRC4 discs per speck is higher by a factor of 1.65 on average (Fig. 5C). Consequently, we found that the density of NLRC4 discs in a speck varied by a factor of around 8 (Fig. 5D). Imaging of NLRC4 and NLRP3 in parallel within the specks, revealed that NLRP3 agglomerates were interspersed with NLRC4 agglomerates. For the NLRP3 agglomerates a diameter of around 50 nm was determined. This is consistent with their in vitro size^50^ and suggests that the NLRP3 agglomerates also represent NLRP3 discs (Fig. 5E). Thus by MINFLUX nanoscopy, we for the first time gained an understanding of the number of NLRC4 discs present in a single speck, as well as their organisation in relation to NLRP3 discs and ASC filaments.

We also wanted to find out whether MINFLUX nanoscopy enables us to demonstrate that hNAIP and NLRC4 colocalise and, if so, to what extent. To investigate this, we imaged hNAIP and eGFP-NLRC4 in parallel within the specks using an anti-hNAIP primary antibody and a secondary nanobody, as well as a GFP-binding nanobody. In line with the patchy localisation of hNAIP in specks, as revealed by STED microscopy (Fig. 4A), we observed far fewer hNAIP localisations than NLRC4 agglomerates (Fig. 5F). To estimate which hNAIP- and NLRC4 localisations may be part of the same inflammasome disc, we took into account the offsets of the antibodies and nanobodies of around 18 nm, a MINFLUX localisation precision of 5 nm and the size of an NLRC4 disc determined in vitro of 32 nm^34,51^, amounting altogether to a maximum of 55 nm (Fig. 5G). We concluded that NLRC4 and hNAIP localisations separated by 55 nm or less are potentially within the same disc. On this basis our analysis indicates that, on average, around 30% of the detected hNAIP molecules were likely located within an NLRC4 disc (Fig. 5H). Taken together, the data provided by MINFLUX nanoscopy suggest that only a small proportion of NLRC4 inflammasome discs within an ASC speck contain hNAIP.

### Nanometre-scale localisation of hNAIP at a disrupted phagosome

We next aimed to visualise the localisation of hNAIP in relation to eGFP-galectin-3 at disrupted phagosomes at the nanometre scale. To this end, WAC(pT3SS)-infected eGFP-galectin-3 expressing macrophages were stained for hNAIP with primary antibodies and secondary nanobodies and imaged with MINFLUX nanoscopy. In line with the patchy localisation of hNAIP at bacteria, as revealed by confocal microscopy (Fig. 2B), we observed hNAIP localisation at distinct sites of the disrupted phagosome in vicinity of eGFP-galectin-3 (Fig. 5I). A 3D reconstruction also revealed that the localisations of individual hNAIPs were inside the phagosome, where bacterial components may be encountered (Fig. 5I). Taken together, these MINFLUX nanoscopy images confirm that hNAIP is recruited to disrupted phagosomes and suggest that hNAIP finds its ligands directly in the vicinity of the bacteria.

## Discussion

During an infection, pathogenic bacteria can activate both canonical and non-canonical inflammasomes in macrophages, resulting in the release of cytokines and pyroptosis^5,6,52^. The activation of inflammasomes can be triggered by a variety of bacterial components and activities^18–23,52^. *Yersinia* taken up by macrophages can activate non-canonical inflammasomes by disrupting the phagosomal membranes that enclose them^13,53^. The current notion is that the NAIPs, the sensors and activators of the canonical NLRC4 inflammasome, bind to the needle and inner rod proteins of the bacterial T3SS in the cytoplasm of infected cells^16,17,33^. However, it has been unclear how proteins such as the needle proteins, which are usually attached to the T3SS, could become accessible to cytosolic immune sensors if they were located within a phagosome in macrophages. Furthermore, the spatial and temporal characteristics of the processes that occur in cells between the sensing of T3SS components by NAIP and the accumulation of NLRC4 inflammasomes in specks — dense condensates of multiple inflammasomes and other components that lead to pyroptosis — remains unknown.

Our study has focused on visualising the process by which the human NLRC4 inflammasome is activated and then further organised in primary human macrophages after uptake of an engineered *Y. enterocolitica* strain expressing the T3SS but no effectors. The specific goal was to visualise the entire process, at a single-cell level and with high spatiotemporal resolution, from the moment the human NAIP/NLRC4 inflammasome senses the T3SS of *Y. enterocolitica* until it becomes integrated in dense condensates known as specks. For this we employed live-cell-and STED-microscopy, as well as MINFLUX nanoscopy, which can reach a localization precision down to 5–10 nm.

We found that, after the enclosing phagosome membrane was ruptured, hNAIP, but not NLRC4 or ASC, was recruited directly to the bacteria in the phagosome remnants. The direct accessibility of a hNAIP ligand (the needle protein of the T3SS) under these conditions strongly suggests that NLRC4 inflammasome activation leading to pyroptosis is triggered by phagosome disruption in our model system. This is further supported by the fact that after the disruption of a phagosome we always observed the formation of a speck and thereafter we could always observe signs of pyroptosis. The times taken for phagosomes to be disrupted after infection, as well as the times taken for a speck to form after phagosome disruption, were highly variable. This suggests that there is significant heterogeneity in how individual macrophages respond to infection with regard to inflammasome activation. One of the most intriguing results of our study was that disruption of one phagosome, which contains in most cases one bacterium, sufficed to cause the formation of a speck in 59% of all observed cases. The fact that only a limited number of ligands will be available on one bacterium suggests extraordinarily effective mechanisms for the amplification of inflammasome activation. Amplification could potentially be due to hNAIP stimulating multiple NLRC4 inflammasomes, which then coalesce in a speck. This process could be aided by NLRP3 inflammasomes that were activated in parallel. The polymerisation of ASC filaments by NLRP3- and NLRC4 discs, which will activate a large number of caspase-1 molecules, is undoubtedly another important amplification mechanism^9^. Another notable result was that the inflammasome components ASC, NLRC4 and NLRP3 could be found in assemblies of various sizes and compositions in the infected macrophages. From these data, we concluded that ASC is apparently recruited to pre-existing NLRP3/NLRC4 clusters as infection time increases.

STED microscopy data of mature specks, here defined as having an area of around 1 µm² and displaying the parallel presence of ASC, NLRP3, NLRC4 and hNAIP, confirmed earlier data indicating that ASC and NLRC4 form an outer and inner ring, respectively, in the two-dimensional projection of a speck^54^. We can now add to these findings that hNAIP is patchily distributed on these rings, indicating that only some of the NLRC4 inflammasomes contain hNAIP.

Investigating mature specks using MINFLUX nanoscopy, which has a localisation precision of around 5 nm, revealed NLRC4 agglomerates consistent with inflammasome discs^34,51^. This allowed for the first time to quantify the number and density of putative inflammasome discs in mature specks. It was interesting to note that disc density varied considerably between specks in a range of ∼60 to ∼480 per μm^3^. Altogether mature specks contained from 35 to 249 individual NLRC4 inflammasome discs. Additional data indicated a higher prevalence of NLRC4 discs than NLRP3 discs in mature specks under our experimental conditions. This is probably because NLRC4 activation predominates following infection with T3SS-producing yersiniae.

MINFLUX nanoscopy also confirmed that hNAIP was much less abundant than NLRC4 in mature specks and also had a more more localised distribution. When measuring the distances between hNAIP and NLRC4 agglomerates, the data indicated that approximately 30% of hNAIP agglomerates were in close enough proximity to an NLRC4 agglomerate to be considered part of the same inflammasome disc. These data are consistent with in vitro assembled NLRC4 inflammasome discs investigated with cryo-EM. Although hNAIP triggered their formation, it was hardly detected in fully formed discs^34^.

In summary, investigating NLRC4 activation in single *Yersinia*-infected macrophages using a combination of imaging techniques at down to nanometre resolution produced several new and unexpected results that will transform our understanding of how NLRC4 inflammasome activation proceeds in cells.

## Materials and methods

### Ethics statement

Approval for the analysis of anonymized blood donations (WF-015/12) was obtained by the Ethical Committee of the Ärztekammer Hamburg (Germany).

### Cell culture

Human peripheral blood monocytes were isolated from buffy coats as described previously^55^. Cells were cultured in monocyte medium (RPMI1640, 20% autologous serum, 1% penicillin/streptomycin) at 37°C and 5% CO2. The medium was changed twice until cells were differentiated into macrophages 7 days after isolation. Macrophages were used for infection one week after the isolation.

If needed, macrophages were transfected with the Neon Transfection System (Invitrogen / Thermo Fisher Scientific) with 5 µg Plasmid-DNA per 10^6^ cells according to manufacturer’s instructions and infected 4 h after transfection.

### Plasmids

pEGFP-hgalectin-3^56^ was a gift from Tamotsu Yoshimori (Addgene plasmid #73080; http://n2t.net/addgene:73080; RRID:Addgene_73080). mScarlet-galectin-3 was generated by replacing eGFP by mScarlet^29^. pLEX-MCS-ASC-GFP^57^ was a gift from Christian Stehlik (Addgene plasmid # 73957; http://n2t.net/addgene:73957; RRID:Addgene_73957). pEGFP-hNAIP and pEGFP-NLRC4 were generated in this study with Gibson Assembly® (New England Biolabs), according to the manufacturer’s instructions. DNA was amplified from pCS2-6myc-hNAIP and pCS2-flag-NLRC4, both a kind gift from Feng Shao^35^, and inserted in the pEGFP-C1 vector (BD Biosciences Clontech). For the cloning of pEGFP-NLRC4 the pEGFP-C1 was cut at the EcoRI and BglII restriction sites.

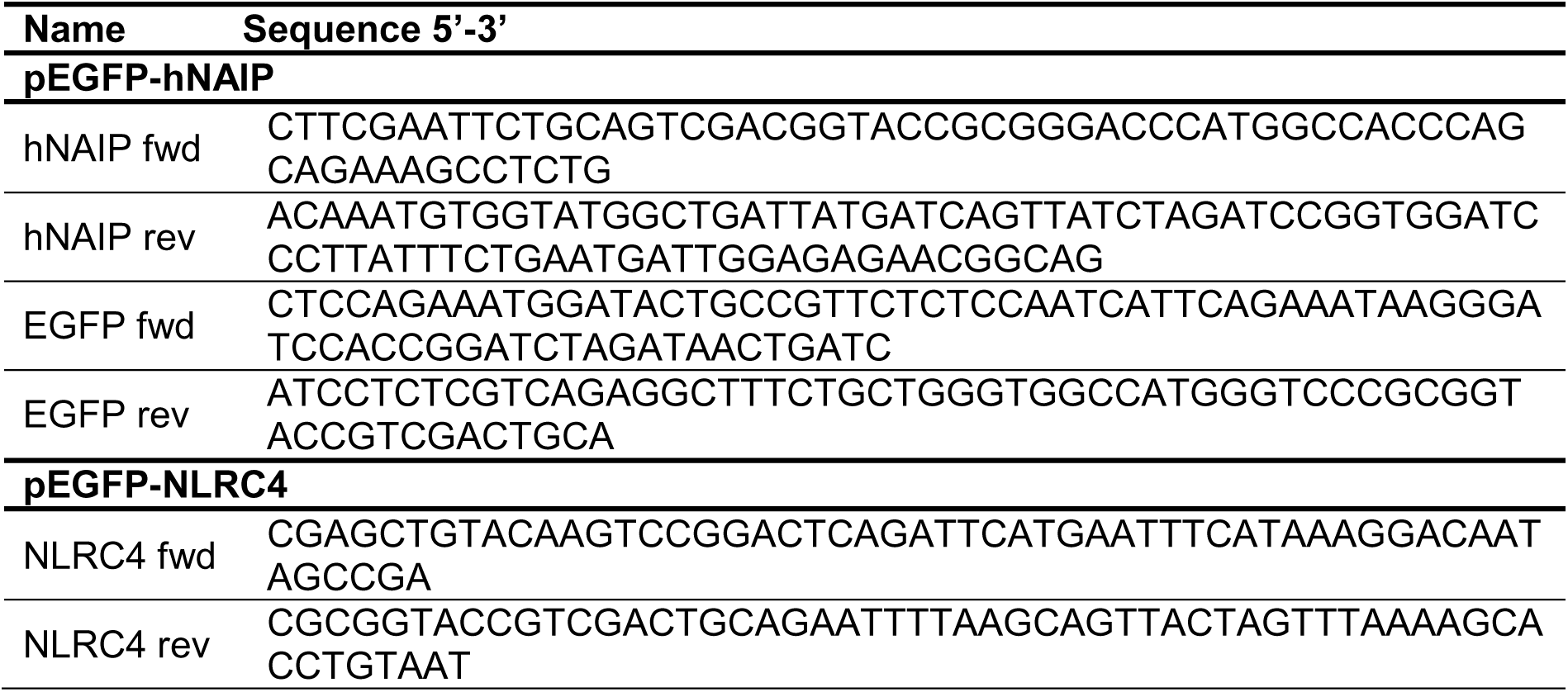

### Bacterial strains

*Y. enterocolitica* strains used in this study derive from the serotype 0:8 strain WA314 harbouring the virulence plasmid pYV08^58^. WAC is the plasmid-less mutant of WA314^58^ and WAC(pT3SS) is a derivative of WAC, which harbours a mini-pYV plasmid encoding for the Type 3 Secretion System (T3SS) and the adhesin yadA^36^.

### Infection of primary human macrophages

*Y. enterocolitica* strains were grown overnight at 27°C and 180 rpm in LB medium containing suitable antibiotics. On the day of infection, bacterial cultures were diluted 1:20 in fresh LB media containing the same antibiotics and incubated for 90 min at 37°C and 180 rpm to induce expression of the virulence plasmid. Bacteria were pelleted at 5000 g, 4°C, 10 min and the pellet was then resuspended in 1 mL ice-cold phosphate-buffered saline (PBS) containing 1 mM MgCl2 and CaCl2 (Sigma-Aldrich, USA). The optical density OD600 was measured and adjusted to 3.6. Cells were then infected using the multiplicity of infection (MOI) indicated for each experiment and incubated at 37°C, 5 % CO2 for the chosen infection time.

### ELISA

Measurement of Interleukin-1β (IL-1β) in the supernatant was performed using the Human IL-1β Uncoated ELISA Kit (Invitrogen / Thermo Fisher Scientific, USA) according to the manufacturer’s instructions, using F96 Maxisorp Nunc-immuno plates (Thermo Fisher Scientific, USA). Measurement was completed with Synergy H1 Multimode Multiplate reader (BioTek /Agilent Technologies, USA). Statistical analysis was performed with GraphPad Prism 11.0.2. Three independent experiments were compared with one-way ANOVA with Tukey’s post-test. P-values ≤ 0.05 were considered statistically significant.

### Live-cell imaging

For live-cell imaging 5×10^4^ hMDMs were seeded in μ-Slide 8 well chamber slides (ibidi, Germany). Cells were placed in the pre-warmed microscopy chamber supplied with 5 % CO2 and infected with *Y. enterocolitica* strains at the indicated MOI. For measurement of cytotoxicity 3 µM DRAQ7TM dye (Thermo Fisher Scientific, USA) was added during infection. SiR-actin (150 nM) was also added during infection. Imaging was performed with the Visitron SD-TRIF mounted to a Nikon Eclipse Tie microscope (Nikon, Japan) with a 63x oil immersion objective (NA 1.40) and the VisiView software (Visitron Systems, Germany) or with the Evident SD mounted to an Olympus IX-83 (Evident Scientific, Japan) with a 20x air objective (NA 0.8) or a 100x oil immersion objective (1.5) and the cellSense software (Evident Scientific, Japan).

### Immunofluorescence staining

For fixed microscopy experiments imaged either with confocal microscopy or STED microscopy 6×10^4^ hMDMs were seeded on glass coverslips (Marienfeld GmbH, Germany) and infected with *Y. enterocolitica* as described above in Infection of primary human macrophages. After required time of infection cells were washed once with PBS and fixed with 4 % PFA (prepared from 16 % PFA; Electron Microscopy Science, USA) in PBS for 10 min. After two washing steps with PBS cells were permeabilized with 0.1 % Triton X-100 in PBS for 15 min at RT. Cells were twice washed with PBS and blocked with 3 % BSA (w/v) in 0.05 % Triton X-100 for 1 h at RT in a humid chamber. Primary antibodies α-ASC (sc51414, Santa Cruz, 1:50-1:100), α-caspase-1 (PA5-17570, Invitrogen, 1:200), α-hNAIP (PA5-115614, Invitrogen, 1:200), α-NLRC4 (#12421, Cell Signalling Technologies, 1:200) and α-NLRP3 (ab4297, abcam, 1:200) were diluted in blocking solution and incubated overnight at 4 °C. Cells were washed five times before incubation with fluorescently labelled secondary antibody, Phalloidin (both 1:200) and DAPI (300nM) for 1 h at RT. After additional 5 times of washing, coverslips were mounted using ProLong™ Glass Antifade Mountant (Thermo Fisher Scientific, USA). All PBS used for immunofluorescence staining contained MgCl2 and CaCl2 (Sigma-Aldrich, USA). Confocal microscopy was performed with the laser scanning microscope FV3000 mounted to an Olympus IX-83 (Evident Scientific, Japan).

### STED microscopy

STED microscopy and corresponding confocal microscopy were carried out in line sequential mode using an Abberior Instruments Expert Line STED microscope mounted to a Nikon Ti-E microscopy body and an Abberior Instruments Mirava STED microscope mounted to an Olympus microscopy body (Abberior Instruments GmbH, Germany). Both were employed for excitation and detection of the fluorescence signal a 60x Plan APO 1.4 oil immersion objective. Pulsed 561, 640 and 680 nm lasers were used for excitation and a pulsed near-infrared laser (775 nm) was used for STED. The detected fluorescence signal was directed through a variable sized pinhole and detected by avalanche photo diodes (APDs) with appropriate filter settings for CF680R, Abberior STAR RED and Abberior STAR ORANGE. Images were recorded with a dwell time between 0.5 and 5 µs and up to 50 line accumulations. The pixel size was set to be 15 nm for 2D-STED, the voxel size was set to be 40×40×50 nm for 3D-STED. The acquisitions were carried out in time gating mode i.e. with a time gating delay of 750 ps and a width of 8 ns. 2D-STED images were acquired with a 2D-STED donut, 3D-STED images were acquired with a 3D-STED donut.

### MINFLUX nanoscopy

For MINFLUX nanoscopy 5×10^4^ hMDMs were seeded in μ-Slide 8 well glass chamber slides (ibidi, Germany). Handling and primary antibody stainings were performed as described above. For MINFLUX imaging primary GFP-nanobodies, secondary rabbit-/mouse-nanobodies or a secondary goat-antibody provided in Abberior or Massive Photonics DNA-PAINT kits were used according to manufacturer’s instructions. MINFLUX samples were incubated with 150 nm goldbeads (Thermo Fisher Scientific, USA) prior to imaging to allow for live sample realignment and MINFLUX beam line monitoring. Single-molecule localization by MINFLUX nanoscopy was enabled by adding adequate concentrations (2-200 pM; depending on the target to be imaged) of the provided imager strands coupled to Abberior DNA-PAINT 660 or Atto 655. MINFLUX nanoscopy, corresponding confocal microscopy and image rendering were carried out using an Abberior MINFLUX setup based on an Olympus IX83 microscopy body equipped with a 100x NA 1.4 oil immersion objective. A 640 nm laser was used for excitation and the iterative MINFLUX localization procedure. A stabilization beam path allowed for live sample realignment by recording a widefield reflection of 150 nm goldbeads caused by illumination with a 980 nm laser. The detected fluorescence signal was directed through a variable sized pinhole (here: 0.83 AU), detected by two APDs detecting different spectral ranges (650-685 nm and 685-720 nm) and later combined. 3D-MINFLUX was performed using the standard 3D-MINFLUX imaging sequence. Multi-color MINFLUX images were acquired using exchange-PAINT. After imaging the first target, imaging buffer containing imager strands binding to the corresponding nanobody of the first target was removed, the sample was carefully washed for at least four times and imaging buffer containing imager strand binding to the nanobody of the second target was added. Imaging of each target was performed for at least 100 minutes. Visualizations and analyses were performed using Abberior Imspector, Paraflux, Imaris and custom MATLAB algorithms for Imaris rendering, quantification and colocalisation analysis.

### Image analyses

#### Confocal and spinning disk microscopy

Images were processed with Fiji/ImageJ (Version 1.54p)^59^. All fixed and live-cell images and movies are depicted as the maximum projection image of the acquired z-stacks, except for Fig. 2B and S1E where only one z-slide is shown.

ASC specks and their components (Fig. 1A, Fig. 3G, H) were counted with the Fiji “Find Maxima” plug-in. Mean fluorescence intensity within single cells was measured to determine ASC-GFP speck formation, phagosome rupture and DRAQ7+ cells. An increase in mean fluorescence was defined as start of each event (Fig. 1C and Fig. 3A, F). Graphs and statistics were created and performed using GraphPad Prism 11.0.2.

#### STED microscopy

STED images were deconvolved using the Abberior TRUESHARP deconvolution. The distance map in Fig. 4B was created by processing binary images of ASC and eGFP-NLRC4 channels in Fiji.

The plot profile in Fig. 4C of the fluorescence intensities of ASC (magenta) and eGFP-NLRC4 (cyan) was measured along the dashed lines indicated in Fig. 4A using Fiji. The arrows indicate how peak to peak measurements were conducted. Each dot in Fig. 4D represents one peak to peak measurement performed on one speck component. For each speck, two measurements per speck component and orientation (xy, xz, yz) using two different line profiles were performed. In total, ASC and eGFP-NLRC4 within five different specks were measured. Statistics were done by using Welch’s t test. Graphs and statistics were created and performed using GraphPad Prism 11.0.2.

#### MINFLUX nanoscopy

MINFLUX localization data were filtered using an effective frequency offset threshold of efo < 100 kHz. Only localization traces with at least 4 localizations were taken into account. Different channels of exchange-PAINT data were aligned based on goldbead localizations derived from MINFLUX beamline monitoring data. For rendering of MINFLUX localization data by Imaris, files were processed using a custom MATLAB script using a pixel size of 4 nm. The voxel size in Imaris was set for 4 × 4 × 2.8 nm (x × y × z) to account for the correction factor of 0.7 of the z-positions when using a #1.5 coverglass with aqueous buffer, as calculated by Gwosch et al. (2020) ^60^. All MINFLUX renderings are depicted using the Imaris surface function (surface detail: 2 nm). For analysing agglomerates as shown in Fig. 5C the localization data of the Imaris surfaces were exported as .xlsx files. A custom MATLAB script was used for quantification of the exported data. If the centre of masses of the surfaces are within a distance of 20 nm they were counted as one. This value was chosen, as the maximal size of the agglomerates is well within the size of single inflammasomes (∼32 nm;)34 including antibody labelling (maximum 2 × 10 nm) and localisation precision (∼5 nm), summing up to a maximal distance of 57 nm (Fig. 5B). The number of NLRC4 inflammasomes per ASC speck is displayed as scatter plot created with Graphpad Prism 11.0.2. Calculations of total NLRC4 inflammasome numbers within a whole speck were calculated based on the assumption that ASC specks are spherical in shape and the layer in the centre of the sphere was imaged. The recorded volume of this spherical layer was calculated and the number of NLRC4 was determined by extrapolating the number of recorded NLRC4 inflammasomes to the volume of a complete ASC speck. Each dot represents the number of NLRC4 inflammasomes within one ASC speck.

To measure the colocalisation ratio of hNAIP to eGFP-NLRC4 data were processed as described above (center of masses of Imaris surfaces are counted as one if they are within a distance of 20 nm). A custom MATLAB script was used to measure the colocalisation between the surfaces of different channels/proteins rendered by Imaris. Surfaces with their centre of masses within 55 nm were defined as colocalising. Considering anti- and nanobody size and the achieved localization precision of ∼ 5 nm, this is the maximal diameter that can be reached by two localizations within one inflammasome (see also Fig. 5G).

## Data Availability statement

All data supporting the findings of this study are available within the article and its supplemental material. Additional materials and data are available from the corresponding author upon reasonable request

## Acknowledgements

The project was funded by the Deutsche Forschungsgemeinschaft (DFG, German Research Foundation) - RTG2771 – project no. 453548970

The laser scanning microscope Olympus FV 3000 and the Evident spinning disk microscope were funded by the DFG (code: INST 152/933-1).

The MINFLUX microscope was funded by the European Regional Development Fund (ERDF) under the Operational Programme Hamburg ERDF 2014-2020, REACT-EU, awarded by the Hamburgische Investitions-und Förderbank (IFB). Grant no. 51164232

We thank the Institute for Transfusion Medicine at the UKE for supply with buffy coats required for monocyte preparation

## Author contribution statement

Susanne Kulnik: Investigation, Methodology, Formal analysis, Writing – original draft, Writing – review & editing

Alexander Westerkamp-Carsten: Investigation, Methodology, Formal analysis, Writing – original draft, Writing – review & editing

Jonas Lübbe: Investigation Shuting Yin: Methodology

Antonio Virgilio Failla: Methodology, Formal analysis

Martin Aepfelbacher: Conceptualization, Funding acquisition, Resources, Supervision, Project administration, Writing – original draft, Writing – review & editing

All authors critically reviewed the manuscript, approved the final version, and agree to be accountable for all aspects of the work.

The authors declare no conflict of interest.

