## Supplemental information for "MINFLUX-nanoscopy of hNAIP/NLRC4 inflammasome activation in single human macrophages"

### Supplementary figure legends

**Figure S1: A)** Increase of eGFP-hNAIP fluorescence during live-cell spinning disk microscopy shown in Fig. 2C. Mean gray values of the area around the bacteria (white box) over time. **B)** Colocalization of eGFP-hNAIP and mScarlet-galectin-3 in infected hMDM after 1 h, MOI100, imaged with a confocal microscope. As also shown in Fig. 2A. Scale bar: 10 µm, Blow up left: 2 µm. **C)** Representative images of endogenous hNAIP staining 120 min after infection of hMDMs with WAC(pT3SS), also shown in Fig. 2B, with parallel staining of DAPI and phalloidin. Scale bar: 10 µm, Blow up: 2 µm.

**Figure S2: A)** Direct comparison of peak to peak measurements of ASC and eGFP-NLRC4 as depicted in Figure 3D STED (4C). Each dot represents the average of two peak to peak measurements performed using two different line profiles on one speck component in one orientation. In total, ASC and eGFP-NLRC4 within five specks were measured from three different orientations. The connection lines between ASC and eGFP-NLRC4 dots indicate measurements of ASC and eGFP-NLRC4 of the same speck. Welch’s t test p-value: 0.0088.

### Supplementary movie legends

**Movie 1: A, B)** Overview and blow-up of live-cell spinning disk imaging of phagosome rupture and eGFP-hNAIP recruitment, according to the stills in Fig. 2C.Scale bars: overview: 10 µm, blow-up: 2 µm.

**Movie 2:** Live-cell spinning disk imaging of phagosome rupture and eGFP-NLRC4 formation as seen to Fig. 2D Scale bar: 10µm.

**Movie 3**: 3D-Reconstruction of speck 1 from Fig. 4A. Depicted is the ASC channel.

**Movie 4:** 3D-rendering of MINFLUX imaging of eGFP-NLRC4 and hNAIP localizations within a macromolecular complex. hMDMs expressed eGFP-NLRC4. eGFP-NLRC4 (cyan) and hNAIP (orange) were imaged consecutively using exchange-PAINT. MINFLUX image: xy-view. Dashed boxes indicate region of blow-ups on the right. Scale bar: 150 nm.

**Movie 5**: 3D-rendering of MINFLUX imaging of hNAIP at the bacteria-containing disrupted phagosome in hMDMs. eGFP-galectin-3 (green) and hNAIP (orange) were localized using imager strands coupled to Abberior DNA-PAINT 660 and proteins were imaged consecutively using exchange-PAINT. Scale bar: 300 nm.

### Supplemental figure S1

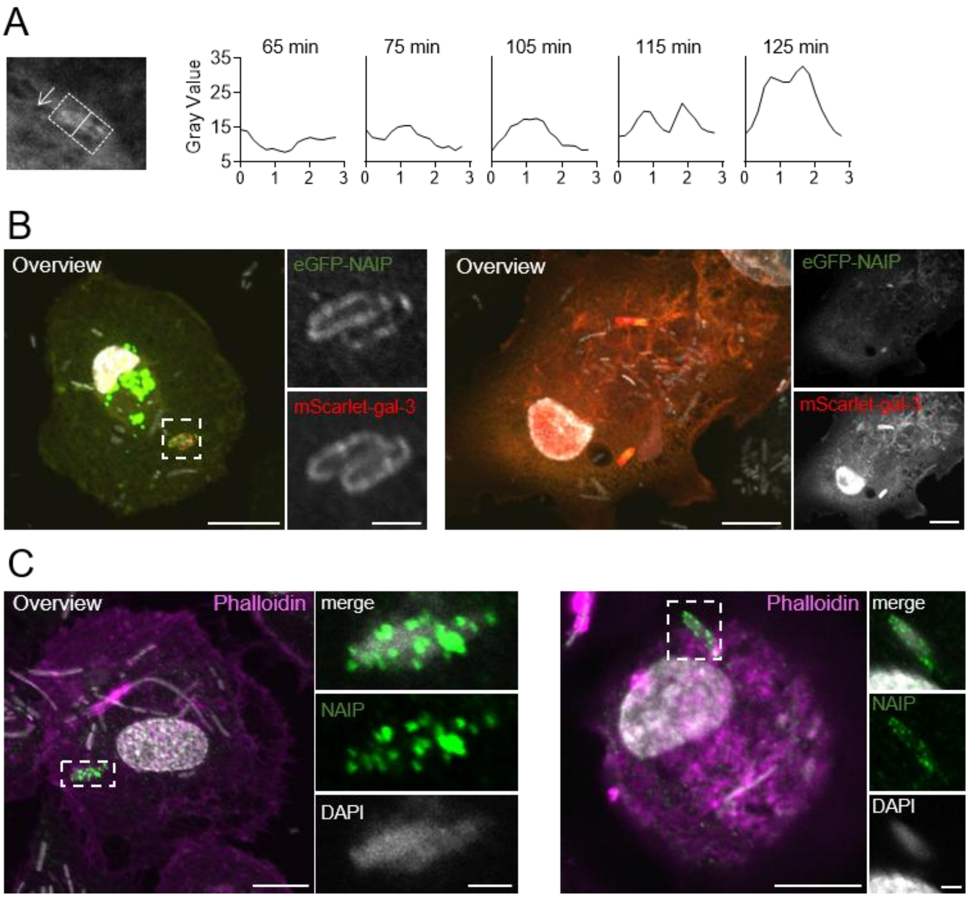


### Supplemental figure S2

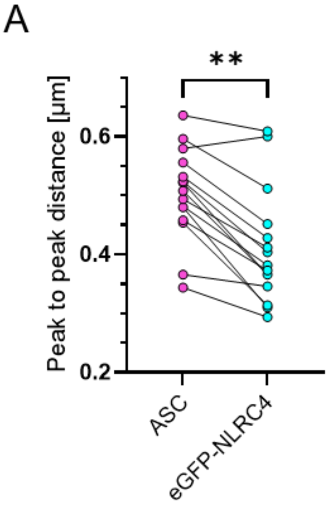
